# Nose-only Exposure to Cherry and Tobacco Flavored E-cigarettes Induced Lung Inflammation in Mice in a Sex-dependent Manner

**DOI:** 10.1101/2022.06.20.496875

**Authors:** Thomas Lamb, Thivanka Muthumalage, Jiries Meehan-Atrash, Irfan Rahman

## Abstract

Flavoring chemicals utilized in electronic nicotine delivery systems (ENDS) have been shown to result in an increase in cellular inflammation, meanwhile, the effects of fruit and tobacco flavors on lung inflammation by nose-only exposures to mice are relatively unknown. We hypothesized that C57BL/6J mice exposed to flavored e-cigarettes would result in an increase in lung inflammation. C57BL/6J mice were exposed to air, propylene glycol/vegetable glycerin (PG/VG), and e-liquids “Apple”, “Cherry”, “Strawberry”, “Wintergreen”, and “Smooth & Mild Tobacco”, for one hour per day for a three day exposure. Quantification of flavoring chemicals was measured by proton nuclear magnetic resonance spectroscopy (^1^H NMR), differential cell counts by flow cytometry, pro-inflammatory cytokines/chemokines by ELISA, and matrix metalloproteinase levels by western blot. Exposure to PG/VG, Apple, and Smooth & Mild Tobacco resulted in an increase in neutrophil cell count in lung bronchoalveolar lavage fluid (BALF). Strawberry exposure increased KC levels in BALF while in lung homogenate KC levels were increased in PG/VG, Cherry, and Smooth & Mild Tobacco exposure. Exposure to PG/VG and Cherry increased IL-6 levels and in all exposed mice there was a male-specific decrease in MCP-1 levels in lung homogenate. Mice exposed to PG/VG, Apple, Cherry, and Wintergreen resulted in an increase in MMP2 levels. Our results indicate that female mice exposed to cherry flavored e-liquids and male mice exposed to tobacco flavored e-liquids resulted in an increase in inflammation, while exposure to mint flavored e-liquids resulted in a decrease in inflammatory cytokine and an increase in tissue repair proteins. This study revealed that flavored-based e-cigarette exposure elicited sex-specific alterations in lung inflammation, with cherry flavors/benzaldehyde eliciting female-specific increases in inflammation. This highlights the toxicity of flavored chemicals and the further need for regulation of flavoring chemicals.

## Introduction

ENDS, also referred to as electronic cigarettes (e-cigarettes), are devices that utilize an atomizer to aerosolize a liquid typically composed of PG, VG, nicotine, and flavoring chemicals at various concentrations (Cao et al. 2021). In 2018, the United States had more than 8000 flavors and 250 e-cigarette brands available on the market (Kaur et al. 2018). In 2018, an estimated 8 million U.S. adults (3.2%) were current e-cigarette users with a high prevalence in young adults, with current e-cigarette users increasing to 4.5% in 2019 (Cornelius et al. 2020; Villarroel et al. 2020).

A majority of e-cigarette users list available flavor choices as their reason for initiation (Soneji et al. 2019). The Population Assessment of Tobacco and Health Study found age-dependent flavor preferences: adolescents have higher affinity fruit flavors than adults (52.8% vs. 30.8%), but a decreased preference for both menthol/mint (10.8% vs 17.9%) and tobacco (5.1% vs. 24.5%) (Schneller et al. 2019). The 2021 National Youth Tobacco Survey also found fruit to be the preferred flavor among middle and high school students (71.6%), with mint and menthol trailing at 30.2% and 28.8% respectively (Park-Lee et al. 2021).

In the United States, current e-cigarette users believe that e-cigarettes are less harmful than traditional cigarettes (Amrock et al. 2015; Huang et al. 2019). Despite the fact that many flavoring chemicals are generally recognized as safe for ingestion, current literature is beginning to show that these compounds may pose health risks to e-cigarette users (Kaur et al. 2018). A recent study demonstrated that ethyl maltol, maltol, ethyl vanillin, and furaneol exhibit cytotoxicity towards lung epithelial cells and mouse neuronal stem cells at concentrations found in e-liquids (Hua et al. 2019). Monocytes treated with maltol, *o*-vanillin, and coumarin, and lung epithelial cells treated with maltol, *o*-vanillin, and diacetyl both released significantly elevated levels of IL-8 (Gerloff et al. 2017; Kaur et al. 2018; Muthumalage et al. 2017). Flavoring chemicals such as maltol and *o*-vanillin have been found in both fruit and tobacco flavored e-liquids (Tierney et al. 2016). Additionally treatments with cinnamaldehyde-containing e-liquids decreased phagocytotic capability of macrophages and neutrophils with concomitant increases in pro-inflammatory cytokine/chemokine secretion in the latter (Clapp et al. 2019). E-cigarette use is also beginning to be associated with lung remodeling and fibrotic-like events along with an increased risk in developing respiratory diseases (Bhatta and Glantz 2020; Hariri et al. 2022; Osei et al. 2020; Wang et al. 2019).

Based on previous literature, with the high preference of flavored e-cigarette use in current users and *in vitro* data showing the induction of an inflammatory response by flavoring chemicals used in e-cigarettes, we hypothesize that nose-only exposure of mice to flavored e-cigarettes would result in lung inflammation. To conduct this study, we exposed mice to five different e-cigarette flavors to 120 puffs daily, a similar number to the daily puffs of e-cigarette users, by utilizing a puffing profile that mimicked the puffing topography of current e-cigarette users and measured pro-inflammatory cytokine levels, BALF cell counts, and lung protease levels to determine lung inflammation (Dautzenberg and Bricard 2015).

## Materials and Methods

### Ethics Statement

Experiments were performed following the standards established by the United States Animal Welfare Act. The Animal Research Committee of the University of Rochester (UCAR) approved the animal experimental protocol conducted at the University of Rochester.

### Animals

Equal number of male and female C57BL/6J mice were ordered from Jackson Laboratory at an age of 6 weeks old. Mice were housed at the University of Rochester for 1 week to acclimatize prior to nose-only tower training. Mice were trained by placing each mouse in the restraints of the Scireq nose-only tower one week prior to e-cigarette exposure. Mice were trained for fifteen minutes on the first day, thirty minutes on the second day, forty-five minutes on the third day, and one hour on the fourth and fifth days.

### E-cigarette Device and Liquids

A Joytech eVIC mini device (SCIREQ, Montreal) with KangerTech 0.15 Ω atomizers/coils (SCIREQ, Montreal) and the Scireq nose-only tower (SCIREQ, Montreal) were utilized for all e-cigarette exposures. E-liquids, PG, and VG were purchased through local vendors/online vendors with e-liquids purchased under the following flavor categories fruit “Apple”, “Cherry”, and “Strawberry”, mint/menthol “Wintergreen”, and tobacco “Smooth & Mild Tobacco”. A 1:1 PG/VG mixture was used for all experiments.

### E-cigarette Exposure

E-cigarette nose-only exposure was performed utilizing the Scireq InExpose system using the Joytech eVIC mini device controlled by the Scireq flexiware software. The puffing profile utilized to expose mice was set at two puffs per minute at an inter puff interval of thirty seconds, with a three second puff duration and a puff volume of 51 ml. Mice were split into seven groups (air, PG/VG, Apple, Cherry, Strawberry, Wintergreen, Smooth & Mild Tobacco) of equal numbers of males (3) and females (3) and exposed using the puffing profile for a total of one hour per day for a three day exposure.

### BALF Collection and Cell Counts

Mice were sacrificed 24 hours after the last e-cigarette exposure and were anesthetized with a mixture of ketamine and xylazine. Mice were lavaged via tracheal catherization three separate times with 0.6 ml of 0.05% fetal bovine serum in 0.9% NaCl. The combined lavage fluids were centrifuged at 2000 rpm for 10 minutes at 4 °C. The supernatant was recovered and stored at -80 °C while the cell pellet was resuspended in 1 ml of 1x phosphate buffer saline (PBS). Total cell counts were measured by staining cells with acridine orange and propidium iodide (AO/PI) and counted using the Nexcelom Cellometer Auto 2000 cell viability counter. Differential cell counts were determined by flow cytometry using guava easyCyte flow cytometer. Cells from BALF were stained with CD16/32 (Tonbo biosciences 70-0161-u500, 1:10) to block nonspecific binding and then cells were stained using a master mix of CD45.1 (Biolegend Cat # 110728, 1:1000), F4/80 (Biolegend Cat # 123110, 1:500), Ly6B.2 (Novus Biological Cat # NBP2-13077, 1:250), CD4 (Invitrogen Cat # 25-0041-82, 1:500), and CD8 (Invitrogen Cat # 17-0081-82, 1:500).

### Protein Extraction

Mouse lung lobes were collected and washed in 1x PBS, blotted dry using a filter pad, and stored at -80 °C. 20 - 30 mg lung tissue were mechanically homogenized in 350 μL of radioimmunoprecipitation assay buffer containing protease inhibitor and EDTA. After mechanical homogenization, samples were placed on ice for forty-five minutes before centrifugation at 14000 rpm for thirty minutes at 4 °C. The supernatant was collected and stored at -80 °C in 50 μL aliquots for ELISA and western blot. To determine the total protein concentration in each sample, the Pierce BCA Protein Assay kit (Thermofisher Scientific, Cat#23225) was used and bovine serum albumin was utilized as the protein standard.

### Pro-Inflammatory Cytokines/Chemokines

Pro-inflammatory cytokine/chemokine keratinocytes-derived chemokine (KC) (R&D DuoSet DY453), interleukin-6 (IL-6) (R&D Duoset DY406), and monocyte chemoattractant protein-1 (MCP-1) (R&D DuoSet DY479) levels were measured using ELISA following manufacturer protocol in BALF and lung homogenate. A dilution of 1:10 was utilized for lung homogenate samples and no dilution was utilized for BALF samples.

### Immunoblot Assay

Equal concentration of lung homogenate samples, 10 μg of protein, were loaded per well of a 26 well 4-15% Criterion Precast Protein Gel (BioRad Cat#5671085) and proteins were ran at 200 V through the gel before being transferred to a nitrocellulose membrane. Non-specific binding was blocked by incubating membranes in 5% non-fat milk in 1x tris-buffer saline with 0.1% tween 20 (TBST) for one hour with rocking at room temperature. Membranes were then probed to determine protein levels using the following antibodies diluted in 5% non-fat milk in 1x TBST: matrix metalloproteinase 9 (MMP9) (Abcam ab38898, 1:1000) and MMP2 (Abcam ab92536, 1:1000) and left rocking overnight at 4 °C. After overnight incubation membranes were washed three times with 1x TBST for ten minutes per wash and then incubated with a secondary goat anti-rabbit antibody (BioRad Cat#1706515, 1:10000) for one hour with rocking at room temperature. Membranes were then washed three times with 1x TBST for ten minutes per wash and signals were measured using an ultra-sensitive enhanced chemiluminescent (Thermofisher Cat#34096) following the manufacturer’s protocol. Images of the membrane were collected utilizing the Bio-Rad ChemiDoc MP Imaging system (Bio-Rad Laboratories). Membranes were then stripped utilizing restore western stripping buffer (Thermofisher Cat#21063) and re-probed for the other MMP and finally for β-actin (cell signaling 12620s 1:2000). Band intensity was determined using densitometry analysis using image lab software and normalized to the levels of β-actin. Fold change in protein levels were relative to the protein levels of air-exposed mice.

### NMR chemical assay

120 μL e-liquids, 600 μL of DMSO-*d*_6_ containing 0.3% tetramethylsilane (Cambridge Isotope Laboratories Inc. Cat#DLM-10TC-25), and 10 μL of a 306 mM solution of 1,2,4,5-tetrachloro-3-nitrobenzene in DMSO-*d*_6_ were combined, then 500 μL of this mixture were introduced into 5 mm Wilmad 528-PP-7 thin wall precision NMR tubes for analysis. ^1^H NMR spectra were acquired on a Bruker Avance 500 MHz NMR spectrometer with 128 scans with a 4.7 s repetition rate, a 30 ° flip angle, with 64k data points. Spectra were processed using Mestrenova with 0.3 Hz line-broadening factor to a final data size of 64k real data points, manually phase-corrected, and baseline corrected using the Bernstein polynomial fit. Flavoring chemical concentrations were determined by comparing the peak integrations of the internal standard to flavoring chemicals, and PG/VG ratio was determined by direct integration of their resonances. All samples were made and ran in triplicate.

### Statistical Analysis

Analysis was preformed using GraphPad Prisma version 8.1.1 utilizing One-Way ANOVA with Dunnett’s multiple comparisons test with data shown as mean ± standard error of the mean (SEM).

## Results

### NMR Analysis of Flavored E-liquids for Flavoring Chemicals

The chemical composition of all e-liquids were assessed by NMR to determine the ratio of PG to VG and quantify key flavoring chemicals in flavored e-liquids. In the Apple e-liquid, the concentration of hexyl acetate was determined to be 0.43 ± 0.04 mg/ml and ethyl maltol was determined to be 0.30 ± 0.05 mg/ml with a 46:54 PG/VG ratio (Table 1). In the Cherry e-liquid, the concentration of benzaldehyde was determined to be 0.12 ± 0.01 mg/ml with a 51:49 PG/VG ratio (Table 1). In the Strawberry e-liquid, the concentration of ethyl maltol was determined to be 0.32 ± 0.05 mg/ml and maltol was determined to be 0.24 ± 0.04 mg/ml with a 50:50 PG/VG ratio (Table 1). In the Wintergreen e-liquid, the concentration of methyl salicylate was determined to be 9.70 ± 0.50 mg/ml with a 49:51 PG/VG ratio (Table 1). Finally, in the Smooth & Mild Tobacco e-liquid, the concentration of maltol was determined to be 1.13 ± 0.02 mg/ml with a 49:51 PG/VG ratio (Table 1).

**Table 1:** Flavoring chemical and propylene glycol and vegetable glycerin quantification in e-liquids. E-liquids were analyzed by H^1^ NMR using a Bruker Advance 500 MHz NMR spectrometer with 128 scans with a 4.7s repetition rate, a 30° flip angle, with 64k data point. Flavoring chemical concentrations and propylene glycol and vegetable glycerin quantification were representative of the average of the three samples ± SEM.

| E-liquids | Flavoring Chemicals | Concentration | PG:VG |
| --- | --- | --- | --- |
| Apple | Hexyl Acetate | $0.43 \pm 0.04$ mg/ml | 46:54 |
| | Ethyl Maltol | $0.30 \pm 0.05$ mg/ml | |
| Cherry | Benzaldehyde | $0.12 \pm 0.01$ mg/ml | 51:49 |
| Strawberry | Ethyl Maltol | $0.32 \pm 0.05$ mg/ml | 50:50 |
| | Maltol | $0.24 \pm 0.04$ mg/ml | |
| Wintergreen | Methyl Salicylate | $9.70 \pm 0.50$ mg/ml | 49:51 |
| Smooth & Mild Tobacco | Maltol | $1.13 \pm 0.02$ mg/ml | 49:51 |

### Alterations in Inflammatory Cell Influx in Lung by Flavors

To determine the effect of flavored e-cigarettes to result in an influx of inflammatory cells, differential cell counts were measured in BALF cells. In all mouse e-cigarette exposure groups, there were no significant alterations in total cell counts or macrophage cell counts compared to air controls (Figure 1A/1B). In combined data, mice exposed to Smooth & Mild Tobacco resulted in a significant increase in the neutrophil cell count compared to air controls (Figure 1C). In male mice, exposure to Smooth & Mild Tobacco resulted in a significant increase in neutrophil cell counts and in female mice, exposure to PG/VG and Apple resulted in a significant increase in neutrophil cell count compared to air controls (Figure 1C). Mice exposed to PG/VG resulted in a significant increase in the CD4 T-cell count compared to air controls (Figure 1D). In all mouse e-cigarette exposure groups, there were no significant alterations in CD8 T-cell count compared to air controls (Figure 1E).

**Figure 1:**
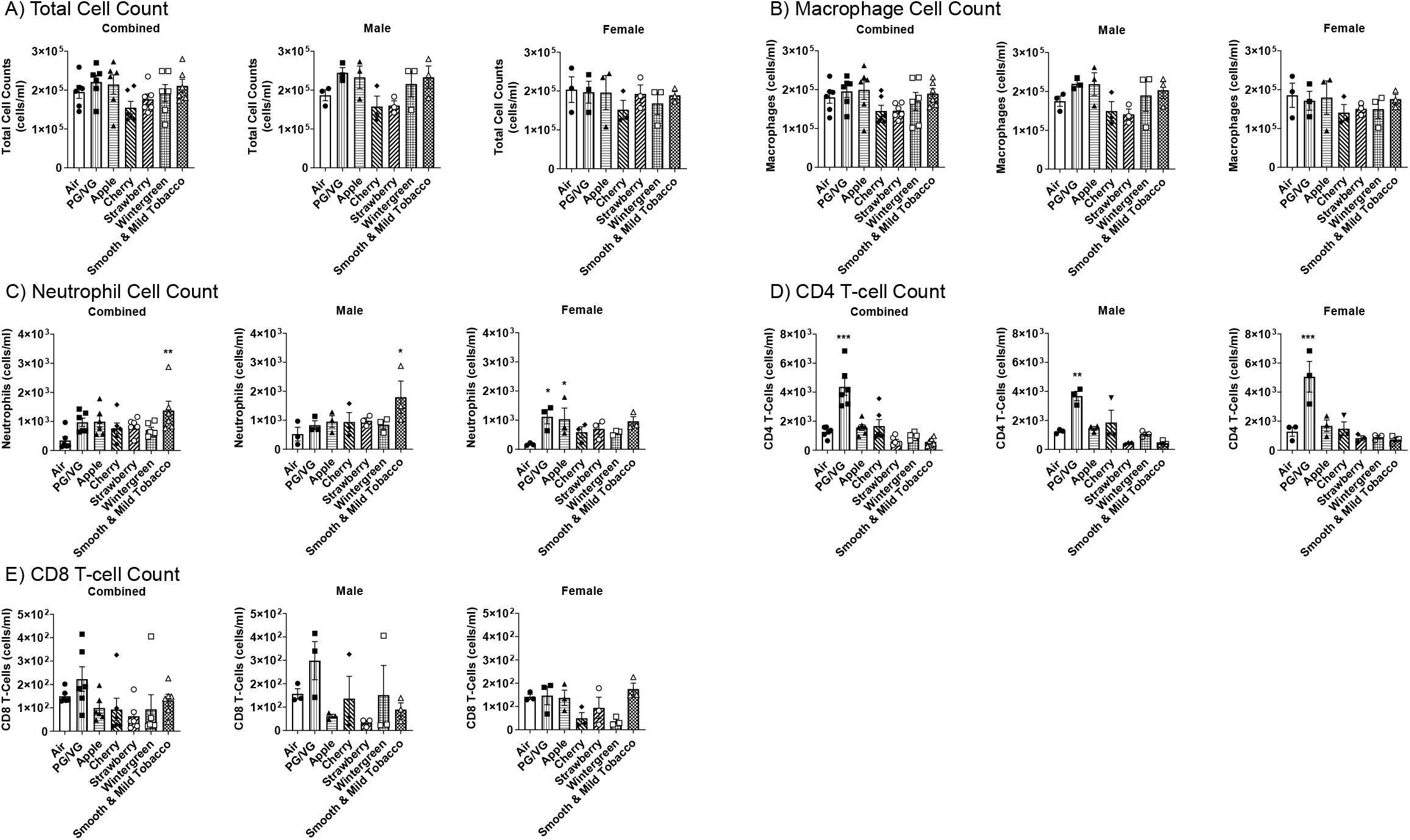
Sex-dependent effects of flavored e-cigarette exposure on inflammatory cell count in bronchoalveolar lavage fluid. Mice were exposed to air, PG/VG, and e-liquid flavors “Apple”, “Cherry”, “Strawberry”, “Wintergreen”, and “Smooth & Mild Tobacco” for 3 days for 1 hour per day. Mice were sacrificed twenty-four hours after the final exposure. (A) Total cell counts were obtained by staining cells with AO/PI and counting with a cellometer. Differential cells were measured using flow cytometry: (B) F4/80+ macrophages, (C) Ly6B.2+ neutrophils, (D) CD4+ T-cells, and (E) CD8+ T-cells. Data are shown as mean ± SEM, with * indicating p < 0.05, ** p < 0.01, and *** p < 0.001 vs air controls. N = 6 for combined groups and N = 3 for male and female only groups.

### Alteration of Pro-Inflammatory Cytokines/Chemokines Levels in Lungs by Flavors

To determine the potential for flavored e-cigarette to elicit an inflammatory response, pro-inflammatory cytokines/chemokines were measured in BALF and lung homogenate. In BALF, KC levels in combined data were significantly increased in Strawberry exposed mice compared to air controls (Figure 2A). In lung homogenate, KC levels in combined data were significantly increased in Cherry and Smooth & Mild Tobacco exposed mice compared to air controls (Figure 3A). In lung homogenate, there was no significant change in any exposed groups in male mice, but, in female mice there was a significant increase in KC levels when exposed to PG/VG and Cherry compared to air controls (Figure 3A). In BALF, IL-6 levels in all exposed mice were not significantly changed compared to air controls (Figure 2B). In lung homogenate, IL-6 levels in combined data were significantly increased in PG/VG and Cherry exposed mice compared to air controls (Figure 3B). In lung homogenate, there was a significant increase in IL-6 levels in male mice exposed to PG/VG as compared to air controls, and female mice exposed to PG/VG and Cherry showed significant increases in IL-6 levels compared to air controls (Figure 3B). In BALF, MCP-1 levels were unchanged in all exposed mice compared to air controls (Figure 2C). In lung homogenate, MCP-1 levels in combined data were significantly decreased in Apple, Strawberry, Wintergreen, and Smooth & Mild Tobacco exposed mice compared to air controls (Figure 3C). In all male mice exposure groups a significant decrease in MCP-1 levels compared to air controls in lung homogenate was observed, whereas for female mice MCP-1 levels were not impacted by the exposures (Figure 3C).

**Figure 2:**
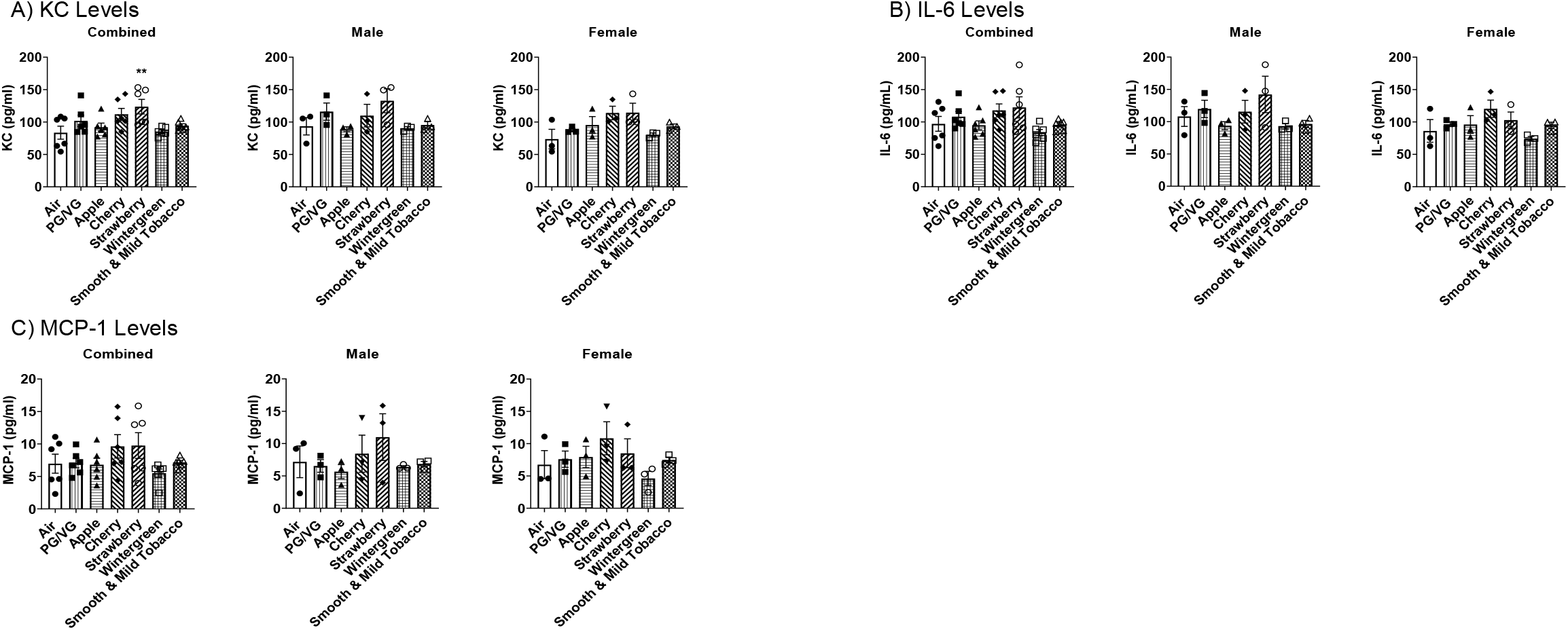
Sex-dependent effects of flavored e-cigarette exposure on pro-inflammatory cytokines/chemokine release in bronchoalveolar lavage fluid. Mice were exposed to air, PG/VG, and e-liquid flavors “Apple”, “Cherry”, “Strawberry”, “Wintergreen”, and “Smooth & Mild Tobacco” for 3 days for 1 hour per day. Mice were sacrificed twenty-four hours after the final exposure. Pro-inflammatory cytokines/chemokines were measured in BALF. (A) KC levels, (B) IL-6 levels, (C) MCP-1 levels. Data are shown as mean ± SEM with * p < 0.05, ** p < 0.01, and *** p < 0.001 vs air controls. N = 6 for combined groups and N = 3 for male and female only groups.

**Figure 3:**
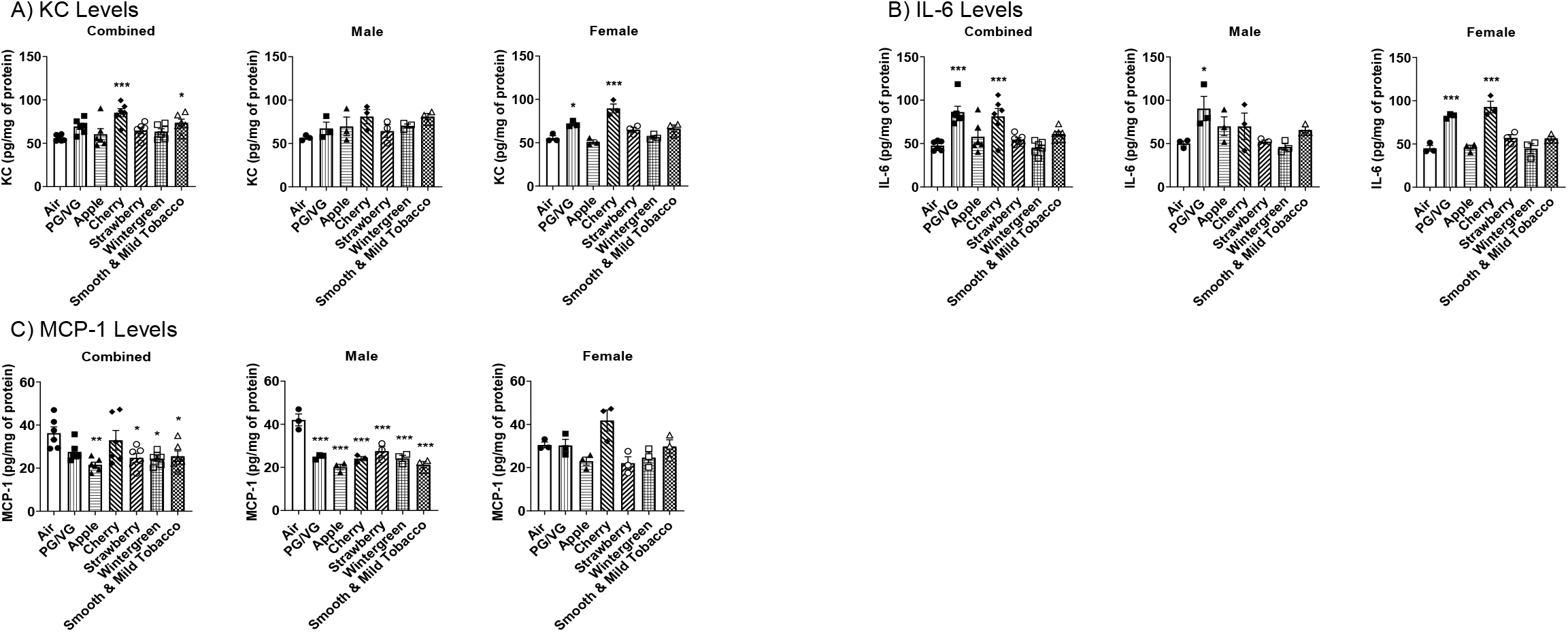
Sex-dependent effects of acute flavored e-cigarette exposure on pro-inflammatory cytokines/chemokine release in lung homogenate. Mice were exposed to air, PG/VG, and e-liquid flavors “Apple”, “Cherry”, “Strawberry”, “Wintergreen”, and “Smooth & Mild Tobacco” for 3 days for 1 hour per day. Mice were sacrificed twenty-four hours after the final exposure. Pro-inflammatory cytokines/chemokines were measured in lung homogenate. (A) KC levels, (B) IL-6 levels, (C) MCP-1 levels. Data are shown as mean ± SEM with * p < 0.05, ** p < 0.01, and *** p < 0.001 vs air controls. N = 6 for combined groups and N = 3 for male and female only groups.

### Alterations in Matrix Metalloproteinase Levels in lungs by Flavors

To determine the effect of flavored e-cigarettes on extracellular remodeling proteins, MMP protein levels were measured in lung homogenate. In all female mice exposure groups, there was no significant change in the relative fold change of MMP9 protein levels compared to air controls (Figure 4B). Exposure to PG/VG, Apple, Cherry, and Wintergreen resulted in a significant increase in the relative fold change of MMP2 protein levels in female mice compared to air controls (Figure 4B). Male mice exposed to Apple displayed a significant decrease in the relative fold change of MMP9 protein levels compared to air controls (Figure 4B). In male mice exposed to PG/VG, Apple, Cherry, and Wintergreen resulted in a significant increase in the relative fold change of MMP2 protein levels compared to air controls (Figure 4B).

**Figure 4:**
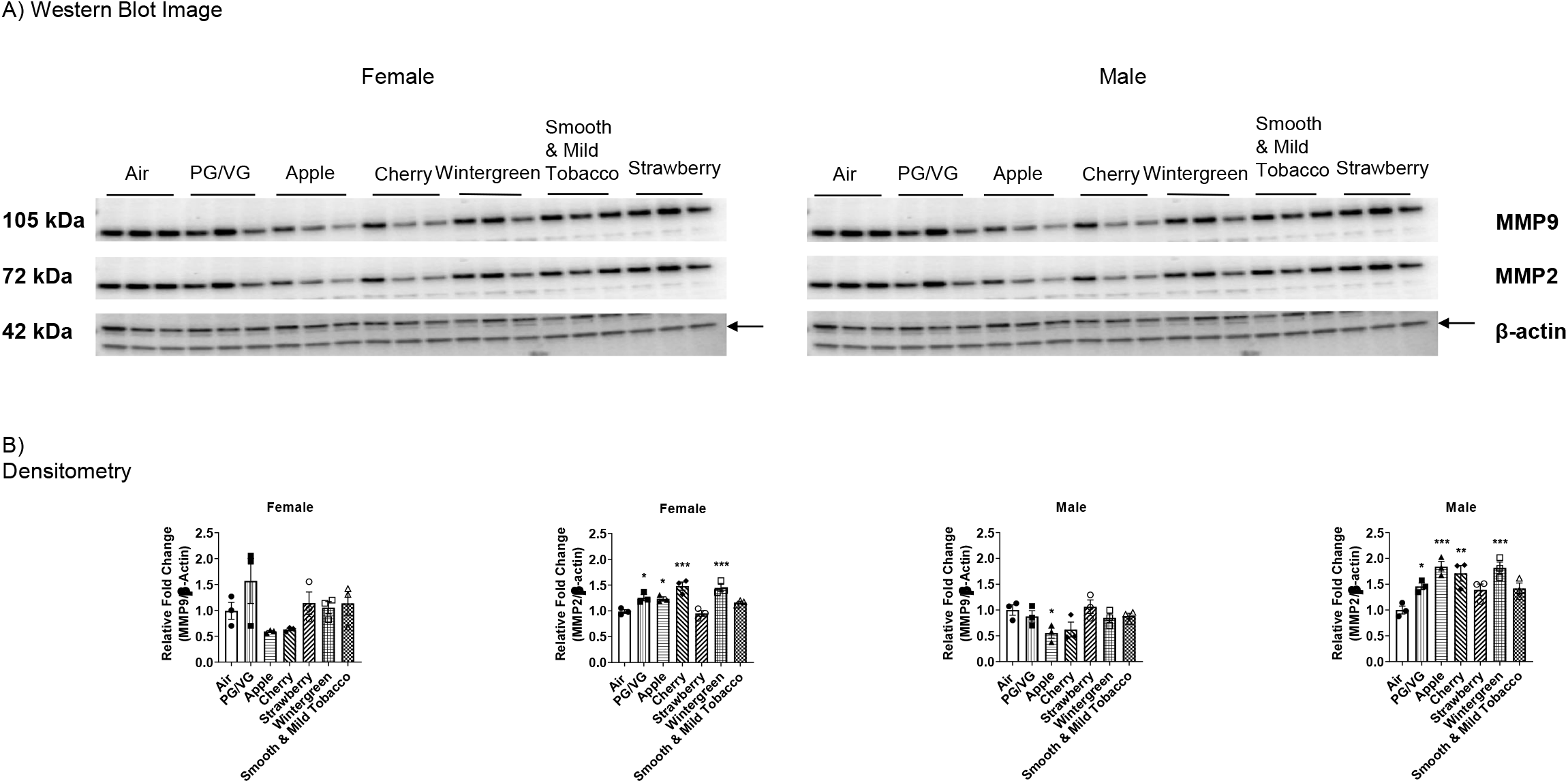
Effects of acute flavored e-cigarette exposure on matrix metalloprotease protein levels in lung homogenate. Mice were exposed to air, PG/VG, and e-liquid flavors “Apple”, “Cherry”, “Strawberry”, “Wintergreen”, and “Smooth & Mild Tobacco” for 3 days for 1 hour per day. Mice were sacrificed twenty-four hours after the final exposure. Protein levels for matrix metalloproteinases were measured in lung homogenate using western blot. (A) Western blot images for MMP2 and MMP9 and loading control β-actin in all male and female exposed mice. (B) Band intensity was measured using densitometry and data are shown as fold change compared to air control mice. Data are shown as mean ± SEM with * p < 0.05, ** p <0.01, and *** p < 0.001 vs air controls. N = 3 for male and female only groups.

## Discussion

In this study, we investigated the immune-inflammatory effects of exposure to flavored e-cigarettes. To determine the potential inhalation effects of flavoring chemicals added into e-liquids, we determined the concentration of five distinct flavoring chemicals (maltol, ethyl maltol, benzaldehyde, methyl salicylate, and hexyl acetate), but the presence of other flavorants are still under investigation. Prior literature also indicates that these compounds have an abundant and widespread presence in market-available e-liquids (Eshraghian and Al-Delaimy 2021; Krusemann et al. 2020; Tierney et al. 2016). While quantification of flavorants are important, a recently published study showcases the inherent variability in lung deposition of flavoring chemicals as a function of inhalation modality: in “lung inhalers” nearly 100% retention of flavorants was observed, but lower retention was observed for “mouth inhalers” (Khachatoorian et al. 2022).

The health effects of ethyl maltol, maltol, and benzaldehyde have been previously studied, but there is currently limited data on the health effects of hexyl acetate and methyl salicylate. Maltol and iron complexes have been found to generate reactive oxygen species (ROS) and can lead to the inactivation of enzymes sensitive to ROS like aconitase (Murakami et al. 2006). Ethyl maltol and iron complexes have been found to be toxic in the liver and kidney of Kunming mice (Li et al. 2017). These interactions with metals are of particular concern in e-liquids due to the presence of metals in the e-liquids within e-cigarette tanks and e-cigarette aerosols (Olmedo et al. 2018). Maltol has also been shown to induce DNA oxidation by enhancing the production of 8-hydroxy-2’-deoxyguanosine (Murakami et al. 2006). Ethyl maltol at a concentration of 12 mg/ml was found to significantly increase ROS production and resulted in a 106% increase in lipid peroxidation (Bitzer et al. 2018).

Benzaldehyde has been found to react with PG to form a benzaldehyde-PG acetal that occurs in higher concentrations in e-liquids containing a greater levels of PG (Erythropel et al. 2019). Both benzaldehyde and benzaldehyde-PG acetal were found to result in a significant decrease in the oxidative burst and phagocytotic capability of human neutrophils (Hickman et al. 2019), and in lung epithelial cells, benzaldehyde-PG acetal exposure resulted in a significant decrease in mitochondrial basal respiration, ATP production, maximum respiration, and spare respiratory capacity (Jabba et al. 2020). Inhalation studies looking into the effects of benzaldehyde have shown that Sprague-Dawley rats exposed to 500, 750, 1000 ppm displayed dose-dependent increases nasal irritation (Andersen 2006), and inflammatory response in C57BL/6J mice (Supplemental Figure 4). This may results in lung inflammatory responses via NF-κB activation and/or mitochondrial dysfunction by flavors/flavoring chemicals, associated with inflammatory mediators release systemically (Lamb et al. 2020; Lerner et al. 2016; Singh et al. 2019).

Similar to total cell count data presented herein, two studies detailing exposures of Balb/c to PG and VG alone and C57BL/6J mice to PG both found no changes in total cell counts (Larcombe et al. 2017; Wang et al. 2020). However, Wang *et al*. found a significant decrease in total cell counts for exposure of PG to C57BL/6J mice (Wang et al. 2019). Similar to the macrophage results detailed herein, exposure to PG or VG and exposure to 70:30 VG:PG with or without vanillin in Balb/c mice, and exposure to PG and exposure to 60:40 PG:VG in C57BL/6J mice did not result in an alteration in macrophage count (Larcombe et al. 2017; Madison et al. 2019; Szafran et al. 2020; Wang et al. 2020). Contrary to the macrophage cell counts from the Wintergreen flavor exposure, Sussan *et al*. showed that C57BL/6J mice exposed to menthol flavor had significant increases in macrophage cell counts (Sussan et al. 2015). In contrast to the results seen in PG/VG, Apple, and Smooth & Mild Tobacco exposures, exposure to PG or VG and exposure to 70:30 VG:PG with or without vanillin in Balb/c mice, and PG exposure and 60:40 PG:VG in C57BL/6J mice did not result in any significant changes in neutrophil count (Larcombe et al. 2017; Madison et al. 2019; Szafran et al. 2020; Wang et al. 2019; Wang et al. 2020). Similar to our Wintergreen flavor exposure, menthol flavor exposure to C57BL/6J mice did not result in any change in neutrophil count (Sussan et al. 2015). In contrast to our CD4 results, C57BL/6J mice exposed to PG resulted in no significant change in CD4 T-cell count (Wang et al. 2019; Wang et al. 2020). Similar to the results herein, there was no change in CD8 T-cells in PG alone C57BL/6J exposure (Wang et al. 2019; Wang et al. 2020).

In line with the increase in IL-6 levels in lung homogenate from PG/VG and Cherry exposures, exposure of C57BL/6J mice to 80:20 PG/VG with 18 mg/ml nicotine found a significant increase in IL-6 RNA levels in the lung tissue (Husari et al. 2016). However, exposure to 60:40 PG/VG in C57BL/6J female mice found no change in IL-6 levels in lung homogenate (Madison et al. 2019). In line with the results from the Wintergreen flavor exposure, menthol flavored C57BL/6J mouse exposure had no change in MCP-1 levels, although this exposure resulted in a significant decrease in IL-6 levels in BALF (Sussan et al. 2015). In contrast to results herein, another C57BL/6J e-cigarette exposure found that PG/VG did not alter IL-6 levels but tobacco flavored exposure significantly increased IL-6 levels in lung homogenate (Glynos et al. 2018). In alternative mice strains ENDS exposure in βENaC resulted in a significant increase in cytokines associated with lung fibrosis (Han et al. 2021). While exposure to PG/VG with nicotine in A/J mice resulted in a significant increase in RNA levels of cytokines associated with chronic obstructive pulmonary disease (COPD) (Garcia-Arcos et al. 2016).

Comparable to results herein, exposure of C57BL/6J mice to PG found that there was no change in MMP9 levels in exposed mice (Wang et al. 2019). However, e-cigarette exposures to PG/VG with nicotine resulted in an increase in MMP9 and other lung protease levels (Garcia-Arcos et al. 2016). Cell studies have found that alveolar macrophages and neutrophils treated with e-cigarette condensate resulted in a significant increase in MMP9 (Higham et al. 2016; Scott et al. 2018). MMP9 levels have also been found to be elevated in the plasma and bronchoalveolar lavage in e-cigarette users (Ghosh et al. 2019; Singh et al. 2019). Matching the MMP2 results herein, increases in MMP2 levels have also been found in mice exposed to PG and increased MMP2 levels have been found in the bronchoalveolar lavage of chronic e-cigarette users (Ghosh et al. 2019; Wang et al. 2019; Wang et al. 2020). Alterations in MMP2 and MMP9 levels due to e-cigarette exposures are important since both MMP2 and MMP9 gelatinolytic activity have been found to be increased in the sputum in both asthmatic and COPD patients (Demedts et al. 2005). MMP2 has also been shown to be upregulated in A459 cells treated with TGF-β indicating a potential role of MMP2 in the abnormal tissue remodeling seen in lung fibrosis (Kasai et al. 2005).

Although the effects of e-cigarettes exposures on cytokine/chemokine levels, MMP levels, and BALF cell counts have different effects in this and other studies, these differences may come down to the methodology for e-cigarette exposures. Each study utilizes different devices and puffing profiles for mouse exposures, along with, different e-liquids with differences in nicotine concentration, flavors, and ratio of PG and VG. These differences between studies showcase the need for a standardized methodology for mouse exposures to reduce potential differences between studies and allow for greater comparisons between studies.

Based on the data collected in this study, flavored e-cigarettes resulted in both increases in lung inflammation and resolution. Mice exposed to PG/VG, Cherry, and Smooth & Mild Tobacco resulted in an increase in lung inflammation due to the increases in KC and IL-6 levels in lung homogenate along with infiltration of neutrophils in BALF. These exposures may also have sex-specific alterations, with Smooth & Mild Tobacco exposure only resulting in a significant increase in neutrophil cell counts in male mice. Meanwhile in Cherry exposure, KC and IL-6 levels were increased in lung homogenate only in female mice. In PG/VG exposures, only female mice had a significant increase in neutrophil cell count and a significant increase in KC levels in lung homogenate. Despite the increases in inflammatory cytokines in Cherry and PG/VG, the increases in MMP2 levels potentially indicate that these exposures have begun to shift away from inflammation and towards tissue repair and resolution. While other exposures such as Wintergreen flavor resulted a decrease in lung inflammation, with a decrease in MCP-1 levels and increases in MMP2 levels. Due to the acute exposure duration, there is still a need for chronic exposures with flavored e-cigarettes to be conducted to determine the potential long-term health effects and the potential sex-specific effects due to flavored e-cigarettes. This study revealed that flavored-based e-cigarette exposure elicited sex-specific alterations in lung inflammation, with cherry flavors/benzaldehyde eliciting female-specific increases in inflammation. This highlights the toxicity of flavored chemicals and the further need for regulation of flavoring chemicals.

## Author Contributions

TL, TM, JMA, IR conceived and designed the experiments

TL conducted the experiments

TL, JMA analyzed the data

TL, TM, JMA, IR wrote and edited the manuscript

