## Supplementary material for "Nose-only Exposure to Cherry and Tobacco Flavored E-cigarettes Induced Lung Inflammation in Mice in a Sex-dependent Manner": Suppl files

Supplementary Figure 1: Full Image of MMP2 for Male and Female Exposed Mice

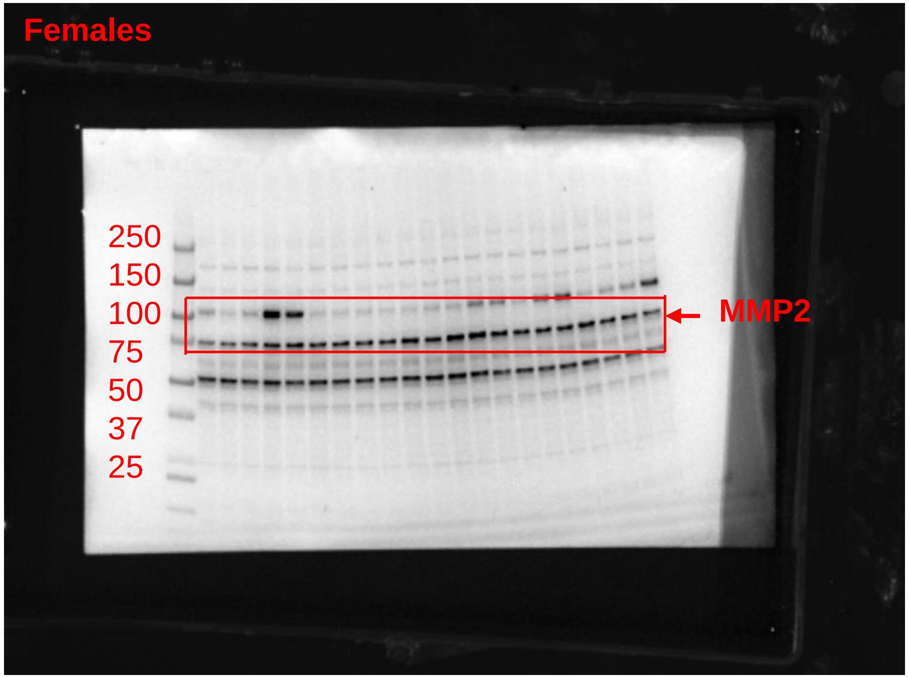


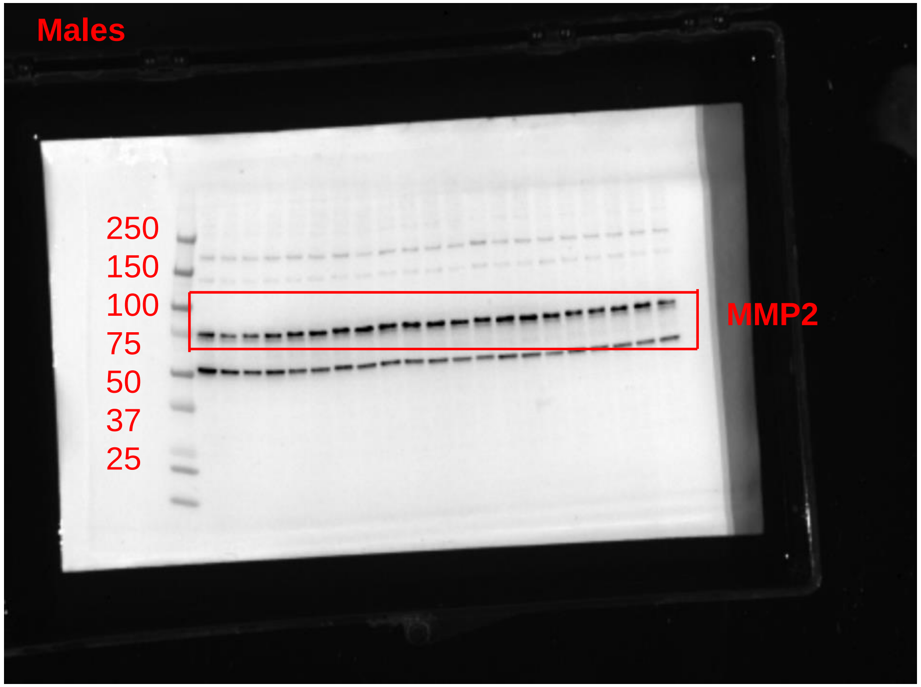


Images represent the full blot for MMP2 with the female mice blot representing the blot after being stripped for MMP9 and re-probed with MMP2 (dilution 1:1000) and with the male mice blot representing the initial probe with MMP2 (dilution 1:1000)

Supplementary Figure 2: Full Image of MMP9 for Male and Female Exposed Mice

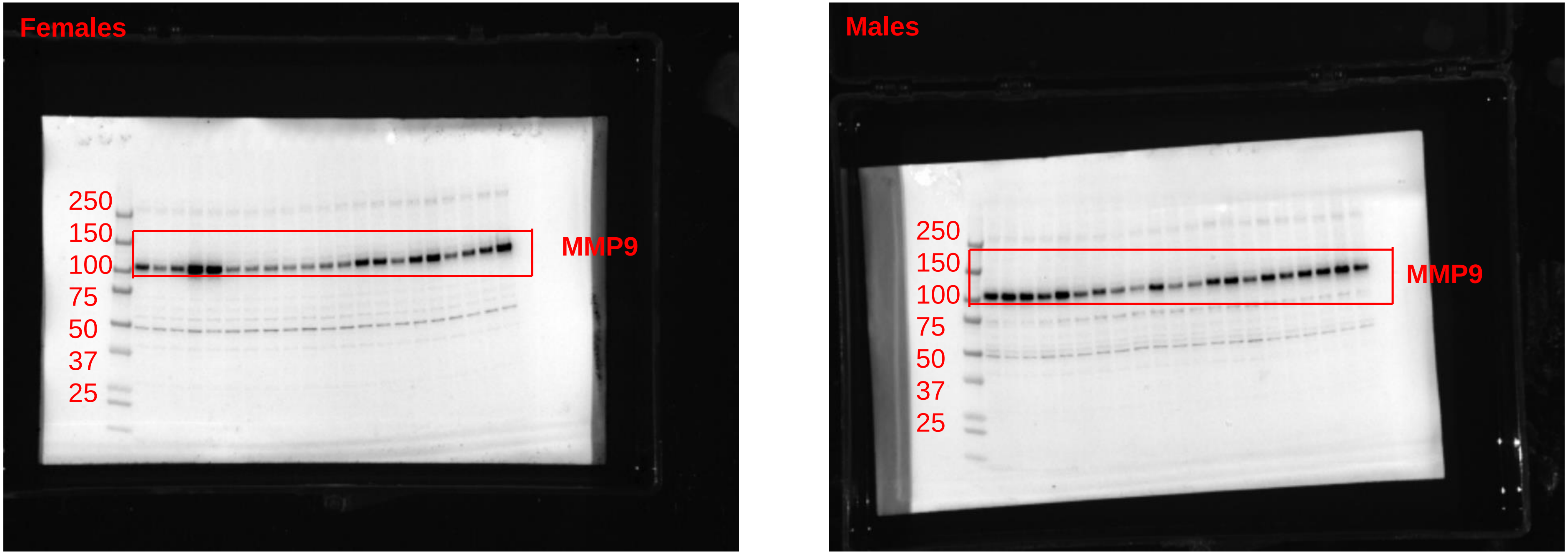


Images represent the full blot for MMP9 with the male mice blot representing the blot after being stripped for MMP2 and re-probed with MMP9 (dilution 1:1000) and with the female mice blot representing the probe with MMP9 (dilution 1:1000)

Supplementary Figure 3: Full Image of β-actin for Male and Female Exposed Mice

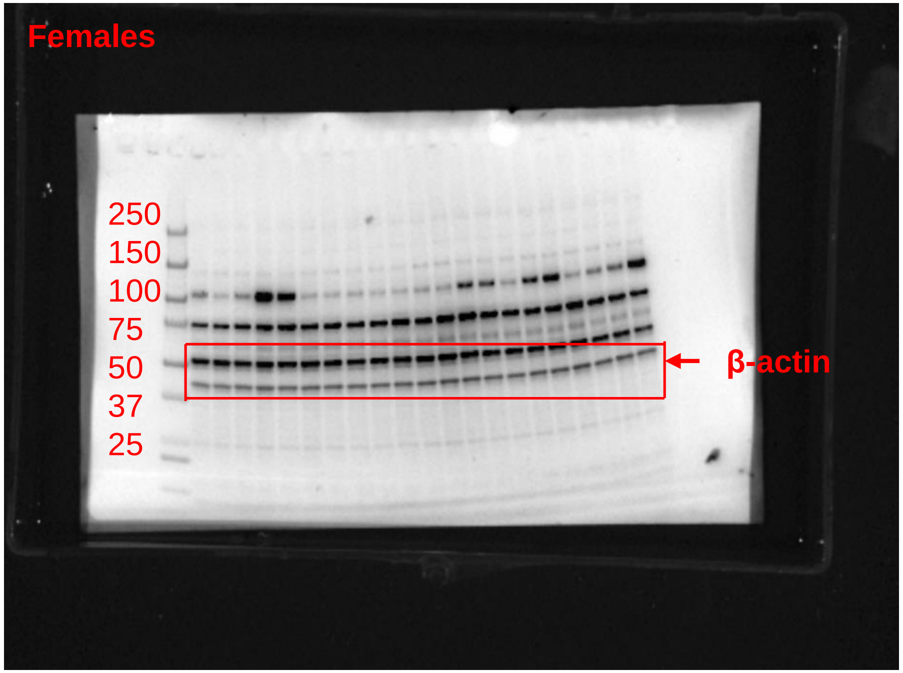


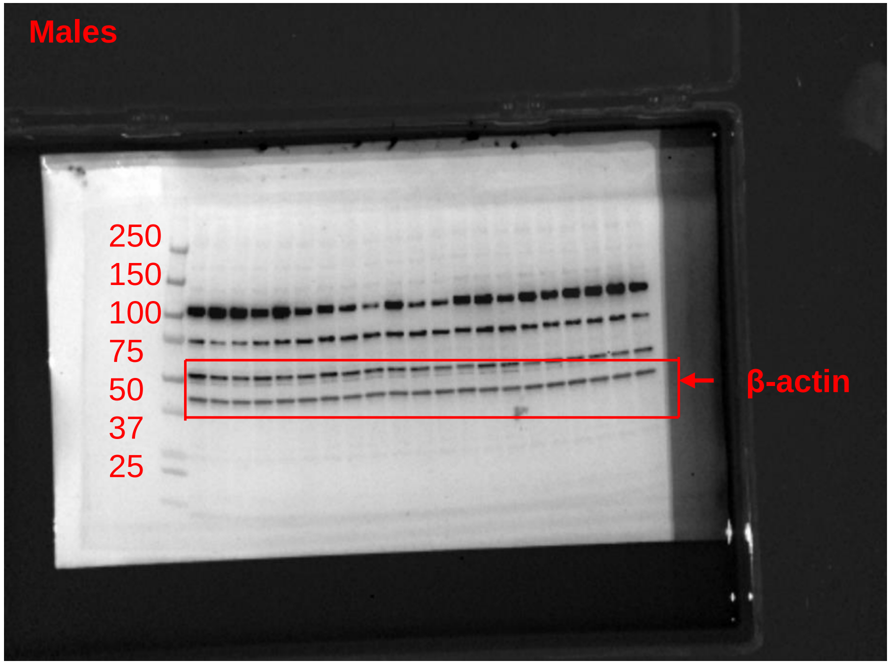


Images represent the full blot for β-actin with both blots representing the image after each membrane has been stripped and finally probed with β-actin (dilution 1:2000)

Supplementary Figure 4: Alterations in pro-inflammatory cytokines/chemokine release in bronchoalveolar lavage fluids due to acute exposure to PG/VG and PG/VG with benzaldehyde

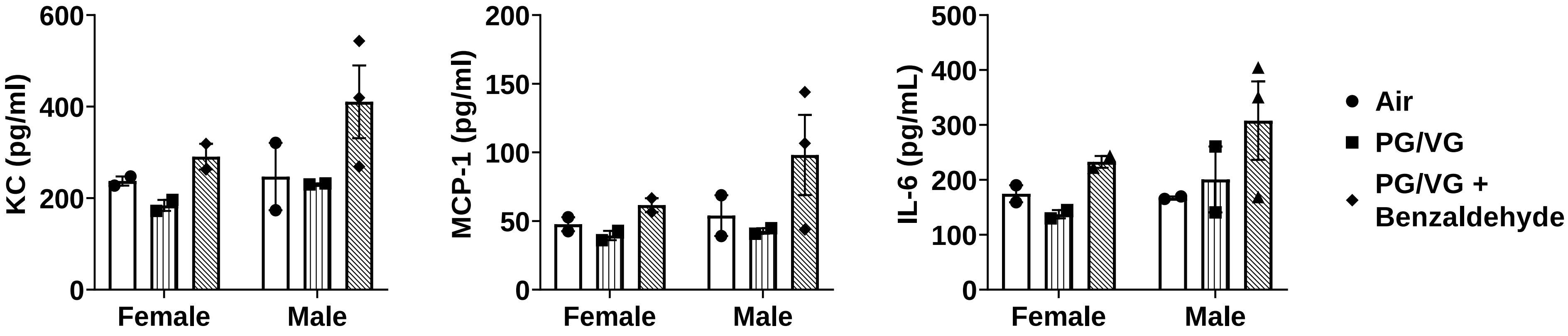


C57BL6/J mice aged, 13-15 weeks old, were exposed for three days to air, 50:50 PG/VG, and 50:50 PG/VG with 280 µg/mL for one hour per day. Mice were exposed to a puffing profile utilized to expose mice was set at two puffs per minute at an inter-puff interval of thirty seconds, with a three-second puff duration and a puff volume of 51 ml. Mice were sacrificed 24-hours post final exposure and were lavaged 3 times with 0.6 ml of 0.05% fetal bovine serum in 0.9% NaCl. BALF was then run for pro-inflammatory cytokines/chemokines KC, MCP-1, and IL-6. Data are shown as mean ± SEM, n = 2-3.
